# *In vitro* and *in vivo* Antiviral Activity of the Acyclic Nucleoside Phosphonate Prodrug LAVR-289 against Poxvirus and African Swine Fever Virus Replication

**DOI:** 10.1101/2025.02.20.634468

**Authors:** Elie Marcheteau, Estelle Mosca, Gaelle Frenois-Veyrat, Sandrine Kappler-Gratias, Laetitia Boutin, Sokunthea Top, Thomas Mathieu, Cyril Colas, Patrick Favetta, Tony Garnier, Peggy Barbe, Mathilde Keck, Daniel Gillet, Alicia Mas, Ali Alejo, Yinyi Yu, Karoly Toth, Getahun Abate, Vincent Roy, Janet Skerry, Kelly S Wetzel, Shamblin D Joshua, Golden W Joseph, Rekha G Panchal, Eric M Mucker, Rajini Mudhasani, Luigi A. Agrofoglio, Frederic Iseni, Franck Gallardo

## Abstract

Poxviruses are double-stranded DNA viruses including relevant zoonotic pathogens with high morbidity. Although African swine fever virus (ASFV) belongs to the *Asfarviridae* family and is not strictly classified as a member of the *Poxviridae*, both fall within the same class of *Pokkesviricetes* that replicate in the cytoplasm, and some poxviruses pose potential biological warfare threats. Among compounds targeting these viruses, acyclic nucleoside phosphonate prodrugs are nucleoside analogues inhibitors of viral DNA polymerases that have been identified as promising agents. However, some limitations related to their toxicity and the rapid emergence of resistance highlight the need for new antiviral molecules. In this study, the new nucleoside analogue LAVR-289 was shown to effectively inhibit the viral replication by intervening early in the viral replication step, targeting a specific domain of the poxvirus DNA polymerase. Using monkeypox virus models, the subcutaneous or oral administration of LAVR-289 demonstrates protective efficacy in infected animal models without toxicity or behavioral modification. The stability *in vivo*, long shelf-life and efficacy make LAVR-289 a promising candidate for further development and stockpiling as a medical countermeasure against dsDNA virus outbreaks. Its broad-spectrum efficacy is a real asset in a context of recurrent viral epidemics, risk of bioterrorism and emergence of resistance strains in the population.

**Highlights:**

- LAVR-289 is a unique acyclic nucleoside phosphonate prodrug targeting viral DNA polymerases.
- LAVR-289 displays antiviral activity against dsDNA viruses, ASFV and poxviruses.
- First report of *in vivo* evaluation of LAVR-289 against MPXV by subcutaneous and oral administration.
- LAVR-289 reduces clinical signs and increase survival in animal models.

## 1. Introduction

The *Poxviridae* family includes viruses that can infect insects (subfamily *Entomopoxvirinae*) and vertebrates (subfamily *Chordopoxvirinae*). These large enveloped viruses, whose double-stranded DNA (dsDNA) genome varies between 128kbp and 360kbp, have an entirely cytoplasmic life cycle^1^. The best-known virus in the *Poxviridae* family is smallpox virus, which has been responsible for terrible, fatal epidemics in humans throughout history until the introduction of systematic vaccination. Following the eradication of smallpox virus, two WHO collaborating centers (in the United States of America and in Russia) preserved samples to pursue international research into i) antiviral agents, ii) improved vaccines, iii) the genetic structure of the virus and iv) the pathogenesis of smallpox^2^. Smallpox virus belongs to the orthopoxvirus genus, which includes other viruses that can also infect humans, such as monkeypox virus (MPXV, causing agent of Mpox disease), cowpox virus (CPXV) and vaccinia virus (VACV). Smallpox was declared eradicated^3^ in 1980 after a WHO-led vaccination campaign, but the end of vaccination and waning immunity have long fueled fears of its possible re-emergence. This was recently exemplified by the accumulation of sporadic outbreaks of zoonotic poxviruses. At the start of 2022, a study warned of a significant rise in MPXV cases in humans since 2000, especially in the Democratic Republic of Congo, where the virus is endemic^4^. The emergence of clade IIb MPXV in the spring of 2022, which quickly spread to every continent and has led to nearly 100,000 cases, was surprising due to its rapid dissemination and swift adaptation to mainly sexual human-to-human transmission. A new variant of clade Ib is currently spreading, with over 18,000 suspected cases and 575 deaths as of this writing^5^. This recent MPXV outbreak serves as a reminder of the high pandemic potential of poxviruses in the human population.

To date, two small molecules (Fig. 1A) have been approved by the FDA for the treatment of smallpox virus^6,7^. One of the drugs, tecovirimat (TPOXX), which targets the viral protein F13 involved in the maturation of viral particles^8^ was shown to be active, *in vitro*, against the MPXV strain circulating during the 2022 outbreak^9^. However, recent NIH reports on two clinical trials (PALM007 and STOMP) performed in order to examine safety and efficacy of tecovirimat during the 2022 outbreak emphasized the critical finding that, despite tecovirimat being safe, it does not improve Mpox resolution and did not shorten the duration of disease relative to supportive care^10,11^. Furthermore, emergence of MPXV resistance to tecovirimat was also observed, particularly in immunocompromised patients^12,13^. The second drug, brincidofovir (BCDV, Tembexa), an acyclic nucleoside phosphonate prodrug, is approved as an orphan treatment specifically for smallpox. It is a pro-drug whose active metabolite, cidofovir (CDV), exhibits broad-spectrum antiviral activity against various DNA viruses^14^. The use of BCDV in patients infected with MPXV has produced side effects that have led to treatment discontinuation^15^.

**Figure 1:**
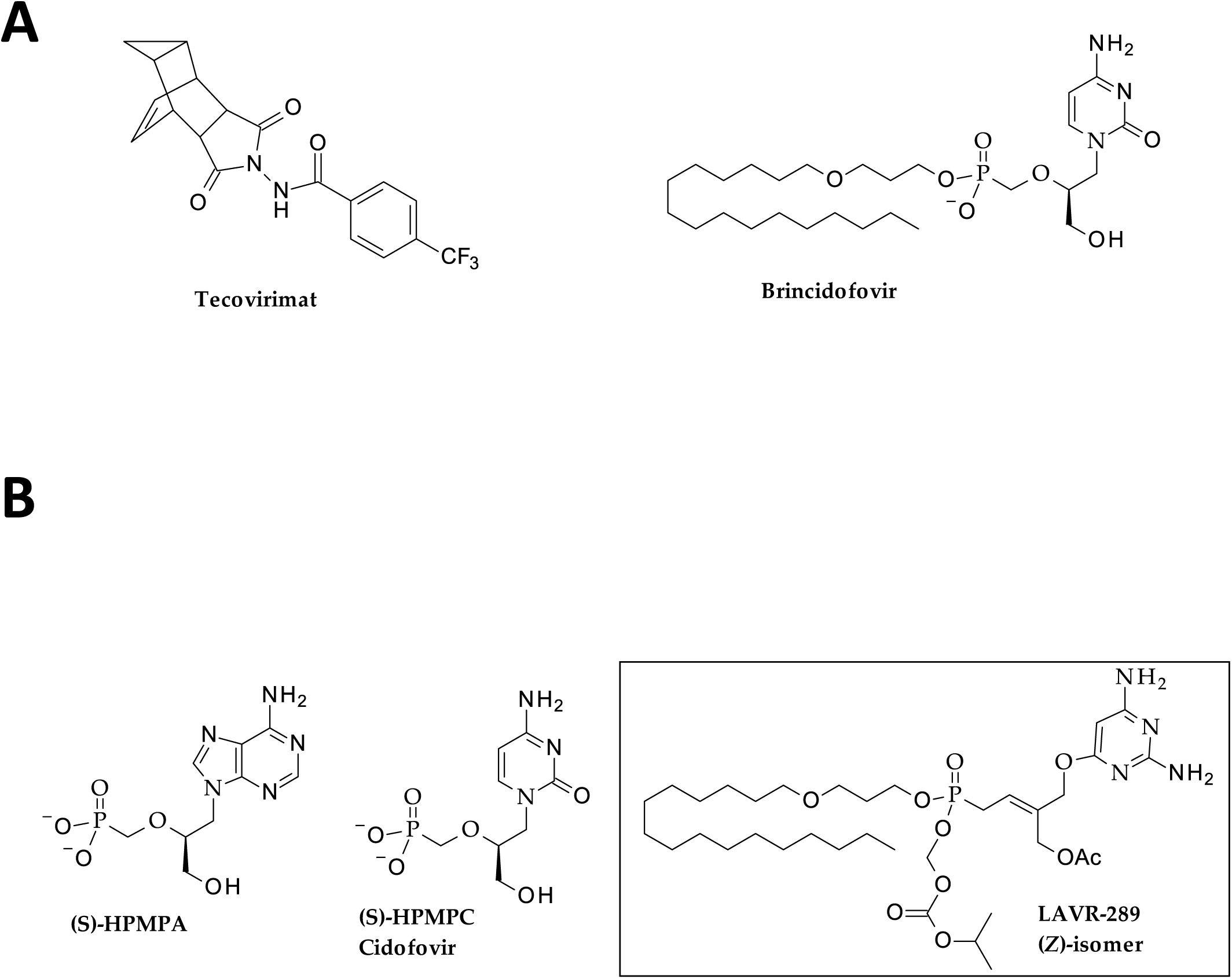
Chemical structure of some active compounds against poxviruses. (A) Tecovirimat and brincidofovir (B) Chemical structure of two acyclic nucleoside phosphonates and LAVR-289.

The African swine fever virus (ASFV) is a dsDNA virus from the *Asfarviridae* family. ASFV is harmless to human but is highly lethal to pigs, leading to severe socio-economic impacts and potential use as a biological weapon for economic terrorism. It requires strict biosecurity measures, including the slaughter of infected animals or those in contact with them to limit the spread of the disease^16^. No vaccine for ASFV has been developed yet, making antiviral agents the best therapeutic option, with acyclic nucleoside phosphonates like (*S*)-HPMPA, (*S*)-HPMPC (CDV) showing promising antiviral activity (Fig. 1B). While (*S*)-HPMPC has demonstrated effectiveness in reducing ASFV replication *in vitro*, it is known to cause nephrotoxicity *in vivo*. BCDV was recently shown to be also partially efficient in preventing AFSV induced mortality in pigs^17^.

Therefore, it is necessary to continue the search for new therapeutics with increased efficiency to enrich the countermeasure arsenal against viruses belonging to both *Poxviridae* and *Asfarviridae* families. We have recently reported the synthesis and *in vitro* antiviral activity of LAVR-289^18^ (Fig. 1B), a new acyclic pyrimidine nucleoside phosphonate prodrug with a 4-(2,4-diaminopyrimidin-6-yl)oxy-but-2-enyl]phosphonic acid skeleton with nanomolar range activity against various DNA viruses (including varicella zoster virus, human cytomegalovirus, human herpes virus and vaccinia virus). Herein, we report on the effectiveness of LAVR-289 to inhibit *in vitro* VACV, MPXV and ASFV in nanomolar range. Moreover, LAVR-289 showed promising efficacy in MPXV infected mouse models and a good safety profile. Interestingly, VACV strains resistant to LAVR-289 were isolated and showed a different mutation pattern to that obtained for CDV.

## 2. Material and methods

### 2.1. Cells and viruses

HeLa (ATCC CCL2), RK13 (ATCC CCL37), Vero (ATCC CCL81), CV-1 (ATCC CCL70) and MM (primary immortalized fibroblast from sheep) cells were used in this study. Cell lines were cultured in Dulbecco’s Modified Eagle’s Medium (DMEM) supplemented with 10 % fetal bovine serum (FBS) (Eurobio-Scientific), 1mM sodium pyruvate (S8636; Sigma Aldrich), L-Glutamine (G7513; Sigma Aldrich) and Penicillin-Streptomycin solution (P0781; Sigma Aldrich). Cells were incubated at 37°C in humidified 5% CO_2_. Vaccinia virus (VACV) TK- RR- (Thymidine Kinase, Ribonucleotide Reductase) TG6002 ANCHOR Santaka^19^ was used for initial testing of LAVR-289. VACV Copenhagen ANCHOR GFP, VACV Copenhagen, CPXV (clinical strain from *Institut de Recherche Biomédicale des Armées,* IRBA, French National Reference Center for Orthopoxvirus) Myxomavirus (MYXV SG33) and lumpy skin disease virus (LSDV) KS1 strain were used to calculate EC_50_ of LAVR-289 and manipulated in biosafety level 2 (BSL-2) laboratory. MPXV Copenhagen and MPXV/France/IRBA2211/2022 were manipulated in BSL-3 (with micro-organism and toxin MOT authorization) at the IRBA. AFSV BA71V ANCHOR and AFSV BA71V were manipulated in BSL-3 at CISA. Clade Ia MPXV (Zaire ’79; V79-I-005) was used for *in vivo* study on CAST/EiJ mice in the United States Army Medical Research Institute of Infectious Diseases (USAMRIID) laboratory. Clade IIb MPXV virus obtained from BEI Resources (NR-2500) was grown in BSC40 cells. Cultures incubated at 37°C were harvested when they exhibited cytopathic effect in at least 75% of the cell layer. Infected BSC40 cell cultures were harvested after freeze-thawing of cell culture flasks. Cultures transferred to centrifuge tubes were sonicated once in a water bath for 30 seconds and pelleted by centrifugation at 3000g. Cell-free supernatants were aliquoted and stored at -70°C. Multiple aliquots were thawed for quantification of virus. Viral titers in culture supernatants were quantified by culturing serial dilutions of the virus in BSC40 cells in Minimum Essential Media (MEM). Handling and storage of clade IIb MPXV cultures were done in Saint Louis University BSL-3 laboratory.

### 2.2. Viral infection of cell cultures and high content imaging

For ANCHOR viruses, cells were plated at a density of 8,000-10,000 cells per well in 96 well plates (Corning Cellbind) in 100µL of complete DMEM. Twenty-four hours post seeding, virus was added to the medium at the indicated MOI. At the indicated time post infection, medium was removed and cells were fixed using formalin (Sigma) for 15 min at room temperature, washed in PBS, stained with Hoechst 33342 (1µg/mL) and imaged using a Thermo CellInsight CX7 high content screening microscope. High content imaging and quantification was described previously^20,21^. Compartmental analysis was used to calculate the total number of cells (number of cell nuclei stained with Hoechst 33342), infection rate (number of ANCHOR GFP positive cells over the total number of cells) and replication level (integrated intensity of the replication center in infected cells using a spot detector algorithm), as described previously^20^.

### 2.3. Antiviral compounds

LAVR-289 was synthetized at a purity of 98% (purity determined by HPLC area) according to the procedure described previously^18^. A 10mM stock solution was prepared in DMSO and stored at -20°C before use for *in vitro* testing. For *in vivo* testing, at 100 mg/kg dosing, LAVR- 289 was dissolved in a solvent composed of 10% DMSO, 10% Cremophor, and 80% PBS at a concentration of 10 mg/mL. For the 50 mg/kg dosing, the compound was diluted in the solvent at a 1:2 ratio. LAVR-289 was prepared on Day 0 and stored at 4°C throughout the study. Prior to daily administration, the compound was allowed to reach room temperature, vortexed to resuspend, and sonicated for approximately 5 minutes. Just before dosing, the suspension was sonicated again. The control article, tecovirimat (from SIGA Technologies Inc.) manipulated during the clade Ia MPXV challenge at USAMRIID, was suspended in a solvent composed of 0.75% methylcellulose (MC) and 1% Tween-80. The compound was homogenized using an OMNI homogenizer to form a milky white solution, then stored at 4°C and stirred overnight. This solution was stable for 14 days. Prior to daily administration, the solution was warmed to room temperature for 30 minutes and homogenized before loading into gavage syringes.

### 2.4. LAVR-289 plasma stability

Human plasma (sterile GTX73265, 100 mL, lot 822101381LiHep, GenTex, France) stored at -28°C was thawed overnight at 4°C. Twenty-seven Eppendorf centrifuge tubes were filled with 240μL of plasma and incubated at 37°C for 30min with agitation (500rpm using two ThermoMixer). Then, 10μL of LAVR-289 solution (11.14 mM in DMSO) was added to each tube. At each measurement time (0, 0.5, 1, 2, 4, 8, 12, 16, and 24h), 750μL of internal standard solution (1.08 mM octylbenzene in acetonitrile) was added to three samples to precipitate proteins. After homogenization, the samples were centrifuged at 4°C for 10min at 10000rpm. Supernatants were collected and transferred to 1.5 mL vials for HPLC-DAD analysis in triplicate. Samples were analysed using an Agilent 1230 Infinity HPLC system with a Zorbax Bonus RP column (150 x 2.1 mm, 2.7µm particle size). The chromatograph was equipped with a 1260 binary pump (G1311B), a 1260 standard autosampler (G1329B), a 1260 thermostated column compartment (G1316A) and a 1260 diode array detector (G4212B) with a Max-Light cartridge cell (1μL volume, 10mm cell path length). Isocratic elution of water/acetonitrile (28/72, v/v) at 0.3mL/min was performed with a 10µL injection at 40°C. Peak areas for octylbenzene (5.9 min) and LAVR-289 (8.5min) were measured at 267nm. A 2mM LAVR-289 stock solution in DMSO was stable for over a week and serially diluted to obtain, after precipitation step, 20, 10, 5, 2.5, 1.25, 0.66, 0.33 and 0.16µM in injected solution. DMSO spiking was under 5% of matrix volume. A calibration curve showed linearity (r² > 0.997) in the concentration range of 0.16 – 20µM.

### 2.5. Selection of LAVR-289-resistant VACV

Drug-resistant viruses were generated as follows: monolayers of Vero cells in 24-well plates (2.9x10^5^ cells per well) were infected with VACV Copenhagen strain at a MOI of 0.01 and then overlaid with 200µL of DMEM containing 250nM of LAVR-289. Forty-eight hours post-infection, infected cells were freeze/thawed three times and centrifuged at 1200 x g for 10min. One twentieth of the preparation was used to re-infect Vero cells in the presence of increasing concentrations of LAVR-289. After 18 to 22 passages, viruses growing in the presence of 2µM of LAVR-289 were harvested. Virus stocks were tested for their ability to form plaques in the absence and presence of 2µM LAVR-289 which showed that resistant strains had been selected. Clonal isolates were selected by plaque purification from infected cells grown in agarose-containing medium with 2µM LAVR-289. Stocks of plaque-purified LAVR-289-resistant VACV were produced and tittered. Viral genome was extracted and Sanger sequencing of the E9L gene was performed. The E9L sequences from LAVR-289- resistant VACV were aligned to the reference genome of VACV Copenhagen (GenBank M35027.1) together with the E9L sequence obtained from a WT VACV that underwent the same number of passages in the absence of LAVR-289.

### 2.6. Viral Plaque Assay and analysis

Confluent Vero cells seeded in six-well plates were infected with WT VACV and mutant VACV suspension containing ∼ 100 plaque-forming units (PFU). After adsorption for 60 min at 37°C, the medium was removed and the cells overlaid with 0.8 ml DMEM and 2% FBS containing the appropriate concentration of LAVR-289. After 1 h at 37°C, 1.6% carboxymethyl cellulose (VWR) diluted in DMEM and 2% FCS was added. The plates were incubated for three days at 37°C in a 5% CO_2_ incubator. Monolayers were fixed and stained in 3.7% formaldehyde, 0.1% crystal violet and 1.5% methanol. The plaques were counted microscopically and imaged.

### 2.7. Generation of recombinant VACVs resistant to LAVR-289 and CDV

The production of recombinant VACV using the CRISPR/Cas9 mediated homologous recombination was described in Boutin *et al. 2022*^22^. Briefly, 5.10^6^ CV-1 cells were infected with WT VACV at a MOI of 0.02 in DMEM supplemented with 0.5 % v/v FCS for 1 h at 37 °C in 5 % CO2 atmosphere. Cells were then electroporated using the Neon Transfection System (Thermofisher) with 2.25 µg of each plasmid: pCMV-Cas9ΔNLS, a donor vector carrying the mutated E9L gene with the LAVR-289-resistant mutations (C356Y or D377V) or the CDV- resistant mutations (A314T and A684V) and pU6-gRNA (Sigma) encoding gRNA targeting E9L. Transfected cells were then seeded in six-well plates and incubated for three days at 37 °C in 5 % CO2. After a single freeze-thaw cycle, the viral suspension was recovered and diluted in DMEM before infection of Vero cells seeded in six-well plates. Infected cells were overlaid with DMEM supplemented with 10 % FCS and 0.6 % agarose and incubated at 37 °C in 5 % CO2. Three days post-infection, 10 to 20 individual plaques were picked and used to infect Vero cells seeded in 96-well plates for two days. For each clone, the viral DNA was extracted and specific genomic regions of E9L were PCR-amplified using the following primers: E9-F: 5’CTAACAAAGAGCGACGTACAAC3’, E9-R: 5’CTTCCCCAATGTTTGGGATTC3’. PCR amplicons were purified and digested with BspEI restriction enzymes (NEB). Digested products were separated by 0.8 % agarose gel electrophoresis and visualized with ethidium bromide. Mutant viruses were further characterized by Sanger sequencing of the E9L gene to ensure the presence of the expected mutations.

### 2.8. Toxicity of LAVR-289 in vivo

For toxicity testing, five groups of BALB/c mice (Charles River) were used and randomized in groups. Six mice were in the untreated group and four in each LAVR-289 treated groups. LAVR-289 was prepared freshly for each injection. Mice were treated by sub-cutaneous (SC) injection on the flank once a day for 10 days. Mice were monitored for acute signs of toxicity (diarrhea, behavior, distress, fluffing) and weighed every day. On day 10, blood was collected via cardiac puncture from each group, and plasma was recovered for clinical chemistry. Collected plasma samples were analyzed on Piccolo Xpress® (from Abaxis Europe GmbH, Griesheim, Germany) with AmLyte 13 disk to determine parameters displayed in Fig. 4B. Quantitative data were shown as means, with error bars indicating the standard error of the mean (SEM). Comparisons between more than two groups were performed using the Kruskal–Wallis test followed by Dunn’s test. Differences were considered significant if *P* value was < 0.05. Statistical analyses were performed using GraphPad Prism 10.0.2.232 software (GraphPad software Inc, San Diego, CA, USA). This procedure has been approved by the French Ethics Committee (APAFIS #37971- 2022071114514238 v3).

### 2.9. Efficacy of LAVR-289 in clade Ia MPXV infected mice

On Day 0, CAST/EiJ mice were infected via intranasal instillation route with clade Ia MPXV (MPXV strain Zaire 79) at a target dose of 10^5^ PFU. Twenty-four hours after virus exposure, LAVR-289 treatment was initiated and was continued once daily via SC injection of 50 mg/kg or 100 mg/kg (as indicated). Tecovirimat was administered at 100 mg/kg by oral delivery. Mice were monitored and scored for signs of disease and weighed individually for 21 days post virus exposure. Animals with clinical score ≥ 8 were euthanized. The scoring criteria were based on four key parameters: appearance, natural behavior, provoked behavior, and weight loss. Appearance was scored from 0 to 3, with severe signs such as hunched posture and porphyrin staining receiving a score of 3. Natural behavior was assessed on a 0 to 8 scale, with immobile animals scoring 8. Similarly, in provoked behavior, unresponsive or cold-to-touch animals received the highest score of 8. Weight loss was scored up to 4, with a ≥30% reduction earning the maximum score. A cumulative score of 8 or higher indicated moribund status, necessitating euthanasia. Toxicity of repeated 100mg/kg dosing was tested in an unexposed group of 3 animals. An animal in the uninfected LAVR-289 100mg/kg died suddenly without presenting any sign of distress. The cause of death in this animal is unknown. Research was conducted under an Institutional Animal Care and Use Committee (IACUC) approved protocol in compliance with the Animal Welfare Act, Public Health Service Policy on Humane Care and Use of Laboratory Animals, and other federal statutes and regulations relating to animals and experiments involving animals. The facility where this research was conducted is accredited by the AAALAC International and adheres to the principles stated in The Guide for the Care and Use of Laboratory Animals, National Research Council, 2011.

### 2.10. Efficacy of LAVR-289 in reducing clade IIb MPXV virus burden in the lung

The animals were fed with standard rodent chow and acidified water ad libitum. On Day 0, mice were infected with 10^4^ PFU of clade IIb MPXV intranasally in 20 μl of MEM. Starting at 3 days post challenge, some mice received 50 mg/kg of LAVR-289 orally in 10% DMSO, 10% Cremophor and 80% PBS in 100µl volume. Control mice received vehicle only. The administration of the compound or vehicle continued once daily for the duration of the study. The mice were observed and weighed daily. During the study, distressed animals were sacrificed as necessary according to SLU IACUC guidelines and left lungs were harvested for determining tissue virus burden. At 7 days post challenge, all surviving animals were sacrificed, and the same samples were collected. Lung samples were homogenized and MPXV tissue viral titer determined by culturing 10-fold serial dilutions of the virus extract in BSC40 cell cultures. All experiments were performed in accordance with federal regulations and following SLU IBC (#2023-00003) and IACUC (#3142).

## 3. Results

### 3.1. Plasmatic, LogP and pH stability of LAVR-289

LAVR-289 is a [(*Z*)-3-(acetoxymethyl)-4-(2,4-diaminopyrimidin-6-yl)oxy-but-2- enyl]phosphonic acid prodrug bearing three biolabil groups (Fig. 1B), a hexadecyloxypropyl (HDP) and *iso*propyloxycarbonyl methyl (POC) moieties at the phosphorus center and an acyl moiety at the unsaturated side chain. HRMS experiment has shown the rapid entry of this compound inside cells, as within few minutes, LAVR-289 presence (tested at 5 µM) becomes undetectable in cell supernatant (*data not shown*). Putative mechanism of entry implies the rapid transfer of the LAVR-289 across cell membrane due to the HDP moiety (Fig. S1A). The prodrug is likely to be metabolized by cellular enzymes, encompassing phosphodiesterase, carboxyesterase and hydrolase esterase to liberate the phosphonic form intracellularly. Since the order of release for the mono-, di-, and tri-deprotected derivatives is unknown, nine intermediate metabolites of LAVR-289 can be expected (Fig. S1B). Its degradation profile in human plasma was determined to be 6.16 h (Fig. S2A), corresponding to the time from which LAVR-289 lost half of its initial concentration. The kinetic profile follows a first-order kinetic model, showing a plasma stability half-life of several hours. Thus, LAVR-289 has a high likelihood of reaching numerous organs, primarily in its hydrophobic prodrug form (LogP = 7.3), suggesting strong cellular permeability potential. This stability is attributed to the presence of the HDP group, as the *bis*(POC) analogue prodrug of LAVR-289 displays a half-life of around 30 min (*data not shown*). Degradation product detection by UPLC-HRMS shows 7 different LAVR-289 forms (Fig. S2B). Of note, no HDP deprotected LAVR-289 could be detected in the human plasma. LogP calculation of the LAVR-289 along with its mono, di and tri deprotected forms indicates that the solubility of the metabolites shifts from highly lipophilic (LogP = 7.3) to highly hydrophilic (LogP = -2.21) for the unprotected diphosphate derivative. The pH stability of LAVR-289 was assessed (Fig. S2C), revealing that LAVR-289 is highly unstable at basic pH but highly stable at acidic pH, indicating its potential to withstand oral administration [see also Quentin-Froignant et al., same issue].

### 3.2. LAVR-289 inhibits orthopoxvirus replication at nM concentration

The antiviral properties of LAVR-289 were investigated on a collection of poxviruses. In a first round of experiments, VACV TK- RR- (TG6002 ANCHOR^19^) was used. TG6002 ANCHOR massively infected cells in the non-treated condition (Fig. 2A top), however, LAVR-289- treated cells at 100nM showed very little sign of infection (Fig. 2A bottom). High resolution imaging of cells infected with VACV ANCHOR and treated or not with LAVR-289 was used to visualize viral DNA accumulation. In untreated conditions, VACV ANCHOR accumulates as expected in large, crescent shaped structures around cell nuclei. Hundreds of fluorescent spots, corresponding to the genome of VACV can be seen in infected cells (Fig. 2B, top). Presence of 1µM of LAVR-289 strongly reduced number of infected cells (Fig. S3). In the restricted number of infected cells, VACV replication is blocked at the single genome (or at least single spot) stage (Fig. 2B, bottom left), and rarely at the 2-3 genome stage (Fig. 2B, bottom right). These results are consistent with the expected mode of inhibition by LAVR- 289 at the viral genome synthesis stage [see below and Quentin-Froignant et al., same issue].

**Figure 2:**
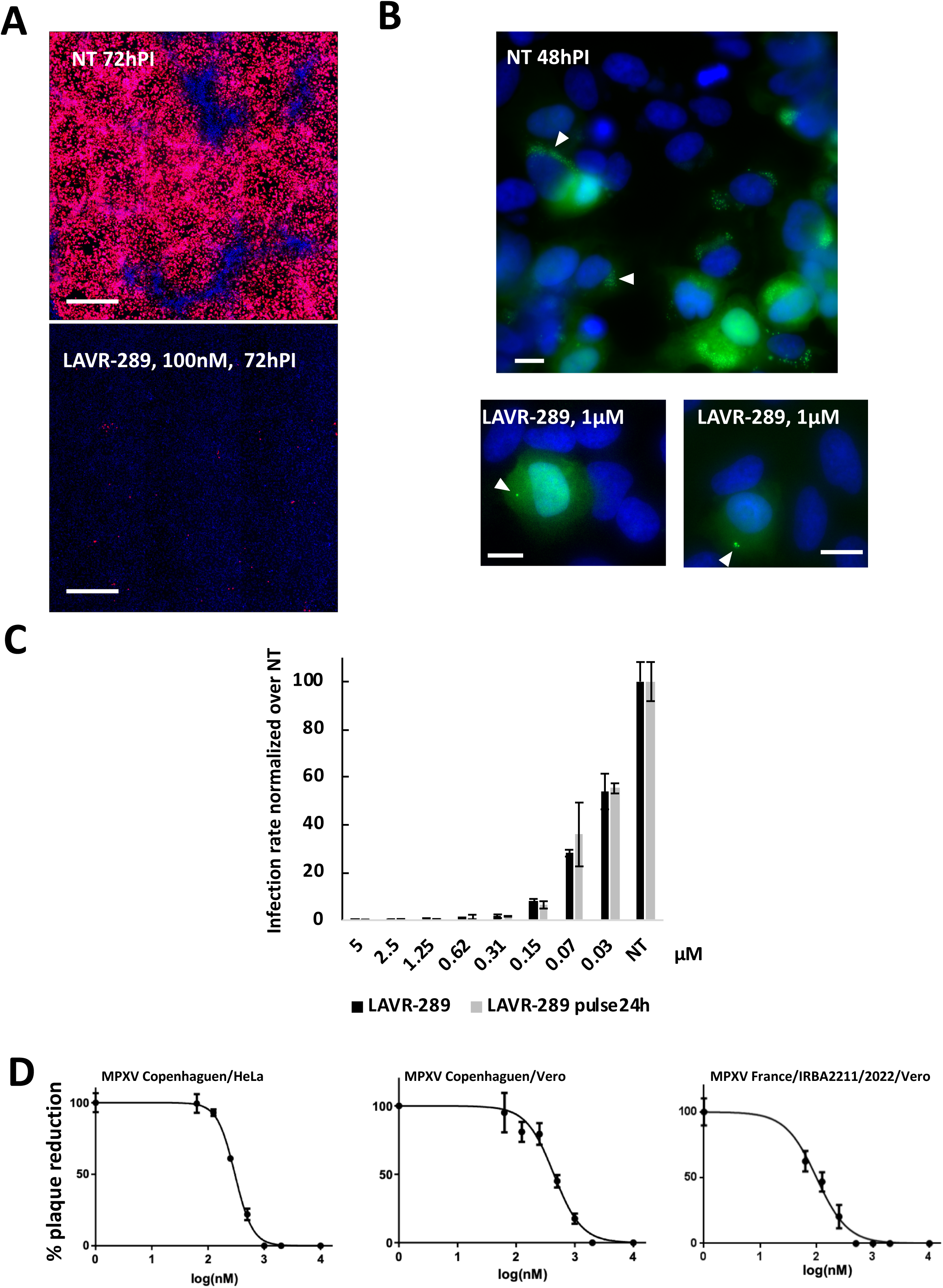
LAVR-289 stably blocks early stage of Vaccinia virus and Monkeypox virus replication at nM concentration. (A) HeLa cells infected with VACV ANCHOR (TG6002, red) at MOI 0.05 and treated or not with LAVR-289 at 100nM and fixed 72h PI. Pictures presented are 25 fields (10X) reconstruction of the cell population, Hoechst 33342 stains nuclei (bar=800µm). (B) High resolution imaging (60X) of cells infected with VACV ANCHOR (Copenhagen strain) at MOI 0.5 and treated or not with LAVR-289 at 1µM and fixed 48 h PI. Arrowheads indicate viral factories (bar=20µm). (C) Percentage of infection normalized over control of LAVR-289 treated cells either continuously or in pulse chase experiment 48 h PI at MOI of 0.5. Dark bars represent continuous treatment and gray bar represent 24 h treatment followed by a wash and release in fresh media for 24 h without LAVR-289. Calculated EC_50_ at 48h in continuous treatment is 40nM. (D) Percentage of plaque obtained on either HeLa (left) or Vero (middle) cells infected with 100 pfu of MPXV (Copenhagen strain), calculated EC_50_ are 300nM and 440nM, respectively. Percentage of plaque observed on Vero cells infected with 100 pfu of MPXV IRBA CNR France 2022 (right). Calculated EC_50_ is 99nM.

To investigate if LAVR-289 is able to stably inhibit VACV infection progression, pulse chase experiments were performed (Fig. 2C). Dose response study shows that at 48 h, LAVR-289 above 150 nM is able to completely prevent VACV replication in cell culture. Calculated EC_50_ in this experiment is 40 nM. Pulse chase experiment does not show any modification of the inhibition capacities, indicating that LAVR-289 activity and/or presence is stable over time and/or that modification of the viral genome is a terminal phenotype and that the LAVR-289 incorporating viral genome cannot be repaired. By performing plaque reduction assays, we further analyzed whether LAVR-289 has a similar antiviral activity on the ancestral MPXV Copenhagen strain and the more recent variant circulating during the 2022 epidemic. LAVR- 289 was tested on the MPXV Copenhagen strain infecting HeLa cells (Fig. 2D left), Vero cells (middle) or Vero cells infected with MPXV/France/IRBA2211/2022 (right). LAVR-289 achieved an EC_50_ of 300 nM, 440 nM and 99 nM, respectively. Combined with toxicity experiments (Fig. S2B), maximal selectivity indexes (SI) reach 593 on Vero cells infected with MPXV/France/IRBA2211/2022. LAVR-289 was also able to inhibit a CPXV (clinical strain EC50 500nM) and other veterinary poxviruses such as MYXV (SG33 strain, EC50 of 30nM) LSDV (KS1 strain, EC50 1.1 µM) (data not shown). Thus, LAVR-289 is highly effective in inhibiting poxviruses replication in cell culture with a very large therapeutic window.

### 3.3. LAVR-289 inhibits replication of the African swine fever virus in vitro

AFSV causes major burden in swine livestock, triggering strong economic consequences and herd culling, and no antiviral or vaccine has been approved to date. We used a recently developed AFSV ANCHOR BA71V strain^23^ to test the antiviral activity of LAVR-289 on this virus. As expected, the AFSV ANCHOR strain readily infects Vero cells and forms large replication centers rapidly after infection and are located close to the nuclei of cells (Fig. S4A, top). Presence of LAVR-289 at 10 µM concentration decreases both the surface and the number of cells presenting a replication center (Fig. S4A bottom). Dose response experiment (Fig. S4B) using HCS quantification shows that starting at 500 nM, LAVR-289 impacts AFSV replication in a dose dependent manner with calculated EC_50_ of 1.1 µM. No significant impact of the compound on cell viability, as measured by flow cytometry, could be detected (NT dead cells: 1.6%, LAVR-289 20 µM: 2.56%, *data no shown*). This result has been confirmed by titration assays using Vero cells, where a LAVR-289 dose dependent reduction in the viral titer can be observed (Fig. S4C). Altogether, these results show that LAVR-289 can inhibit the replication of the AFSV. Structurally, the AFSV replicative polymerase was found to be more similar to the human DNA polymerase delta than to the DNA polymerase of MPXV, which may explain any differences observed in activity^24^.

### 3.4. LAVR-289 targets the VACV DNA polymerase

To confirm that LAVR-289 targets the viral DNA polymerase, we generated LAVR-289- resistant VACV strains. Three independent infected Vero cells cultures were treated with increasing concentration of LAVR-289 (from 250nM to 2 µM) for 18 to 22 passages. As shown in Fig. 3A, resistant viruses were obtained (from each culture) forming plaques similar to WT virus in size and number when not treated with the drug. However, in contrast to WT VACV, 2µM of LAVR-289 had no detectable inhibitory effect on the viruses. Sanger sequencing of the E9L gene from three independent resistant clones per culture showed that all the clones carried a point mutation in the DNA polymerase gene. These mutations are C356Y or D377V/Y (Fig. 3A). In a similar but independent experiment, a resistant VACV carrying a V388F was also isolated (*data not shown*). Viruses C356Y or D377V/Y displayed no fitness defect when compared to the WT virus growth as shown on Fig. 3B. Interestingly, the 3 LAVR-289 resistant mutations are all located in the poxvirus-specific insert 2 of the DNA polymerase (Fig. 3C and D)^25,26^ and are not the same as those described for CDV- resistant viruses namely A314T and A684V (Fig.3C)^22^. As CDV and LAVR-289 are both acyclic nucleoside phosphonates, we wondered whether a virus resistant to CDV was also resistant to the LAVR-289 drug. To do this, we produced two recombinant VACV using the CRISPR/Cas9 technique^22^: one virus carried the CDV-resistant mutations A314T and A684V and another recombinant VACV carried the LAVR-289 resistant mutation D377V. As expected, VACV^D377V^ displayed a LAVR-289 resistant phenotype if compared to the WT virus (Fig. 3E). Surprisingly, the CDV-resistant VACV was also resistant to LAVR-289 drug. Altogether, our results indicate that LAVR-289 targets the insert 2 domain of the VACV DNA polymerase. Interestingly, even if LAVR-289 and CDV act with a similar mode of action, obtaining resistance mutants targeting different residues in different polymerase structures indicates they may interact with the polymerase differently.

**Figure 3:**
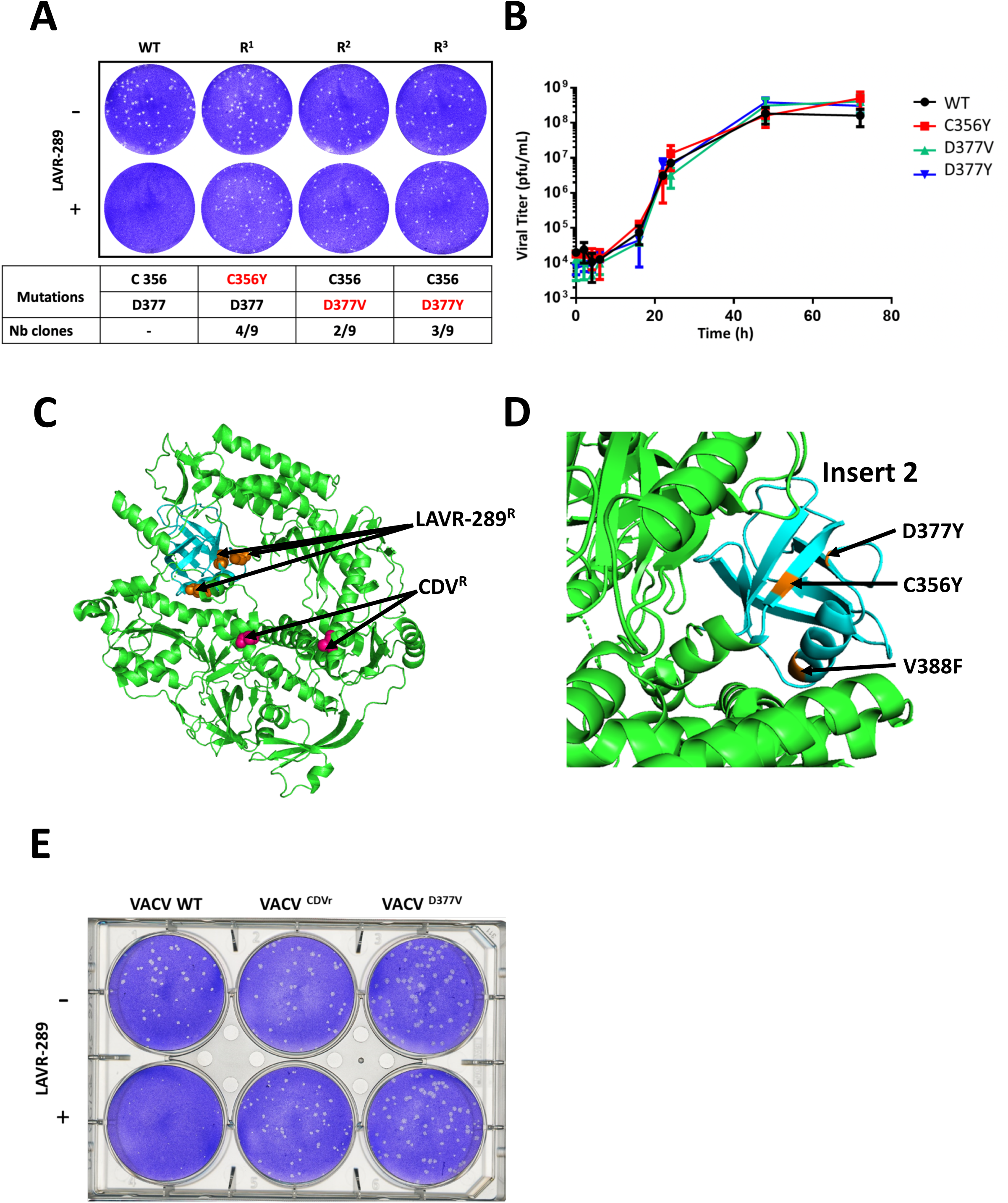
LAVR-289 targets the insert 2 of the poxvirus DNA polymerase. (A) Resistant VACV obtained upon successive passage with increasing concentration of LAVR-289, from 250nM to 2 µM (three replicates R1, R2, R3). Mutated residues are indicated in red. (B) Growth curve of the indicated VACV mutants in Vero cells. (C) 3D structure of the E9 viral DNA polymerase, with position of CDV^R^ and LAVR-289^R^ mutations indicated. (D) Position of the identified LAVR-289 resistance mutations. All mutations are located in the insert 2 of the viral DNA polymerase. C356Y is known to confer resistance to phosphonoactetic acid (PAA). (E) LAVR-289 treatment of Vero cells infected with CDV^R^ and LAVR-289^R^ VACV. CDV^R^ strains is also resistant to LAVR-289.

### 3.5. Subcutaneous administration of LAVR-289 is not toxic in mice

To determine if LAVR-289 displays *in vivo* toxicity, we injected Balb/c mice with indicated concentrations of LAVR-289 administered SC once per day for 10 days. Mice were weighed and signs of toxicity were observed daily (Fig. 4A). At Day 10, no statistically significant differences were determined between controls (vehicle alone) and treated groups, suggesting that the compound is well tolerated in mice. Toxicological effects of LAVR-289 dosing were also investigated from plasma of mice administered with different concentrations of the molecule (Fig. 4B) after 10 days of treatment. Analytes reflecting hepatic, renal, pancreatic, metabolic and muscle parameters, as well as electrolyte were unremarkable. A significant decrease in albumin was observed after administration of 52 mg/kg LAVR-289 (Fig. 4B), but still in range of standards values of the assay (dashed bars). No significant signs of perturbations of other plasma parameters were detected. Pharmacokinetics experiment of LAVR-289 intravenously (IV) injected shows a remarkable half-life of 12,5 hours in mice, compatible with a once a day treatment (Table 1). Maximum tolerated dose (MTD) in mice using dose escalating IV administration showed that the MTD is superior to the maximum tested dose of 100mg/kg (*data not shown*). All together, these results indicate that LAVR-289 is not toxic in mice when administered subcutaneously once a day during a 10-day regimen, can be used at high concentration even in IV and displays pharmacokinetic properties compatible with a once-a-day treatment regimen.

**Table 1:** Pharmacokinetics parameters of LAVR-289 after single 5 mg/kg IV injection in mice. LAVR-289 displays a half-life (T_1/2_) of 12.47 h, a mean resident time (MRT) of 17.50 h, a volume of distribution at steady-state (Vss) of 465.44 L/kg, and clearance (CL) of 443.16 mL/min/kg, indicating a long exposure and slow elimination from the organism. AUC: area under the curve.

| LAVR-289 | t<sub>1/2</sub><br>(h) | C<sub>0</sub><br>(ng/mL) | AUC<sub>last</sub><br>(h*ng/mL) | AUC<sub>inf</sub><br>(h*ng/mL) | AUC/D<br>(h*kg*ng/mL/mg) | AUC Extr<br>(%) | MRT<br>(h) | V<sub>ss</sub><br>(L/kg) | CL<br>(mL/min/kg) |
| --- | --- | --- | --- | --- | --- | --- | --- | --- | --- |
| 5 mg/kg, IV | 12.47 | 74 | 134 | 188 | 37.61 | 28.71 | 17.50 | 465.44 | 443.16 |

**Figure 4:**
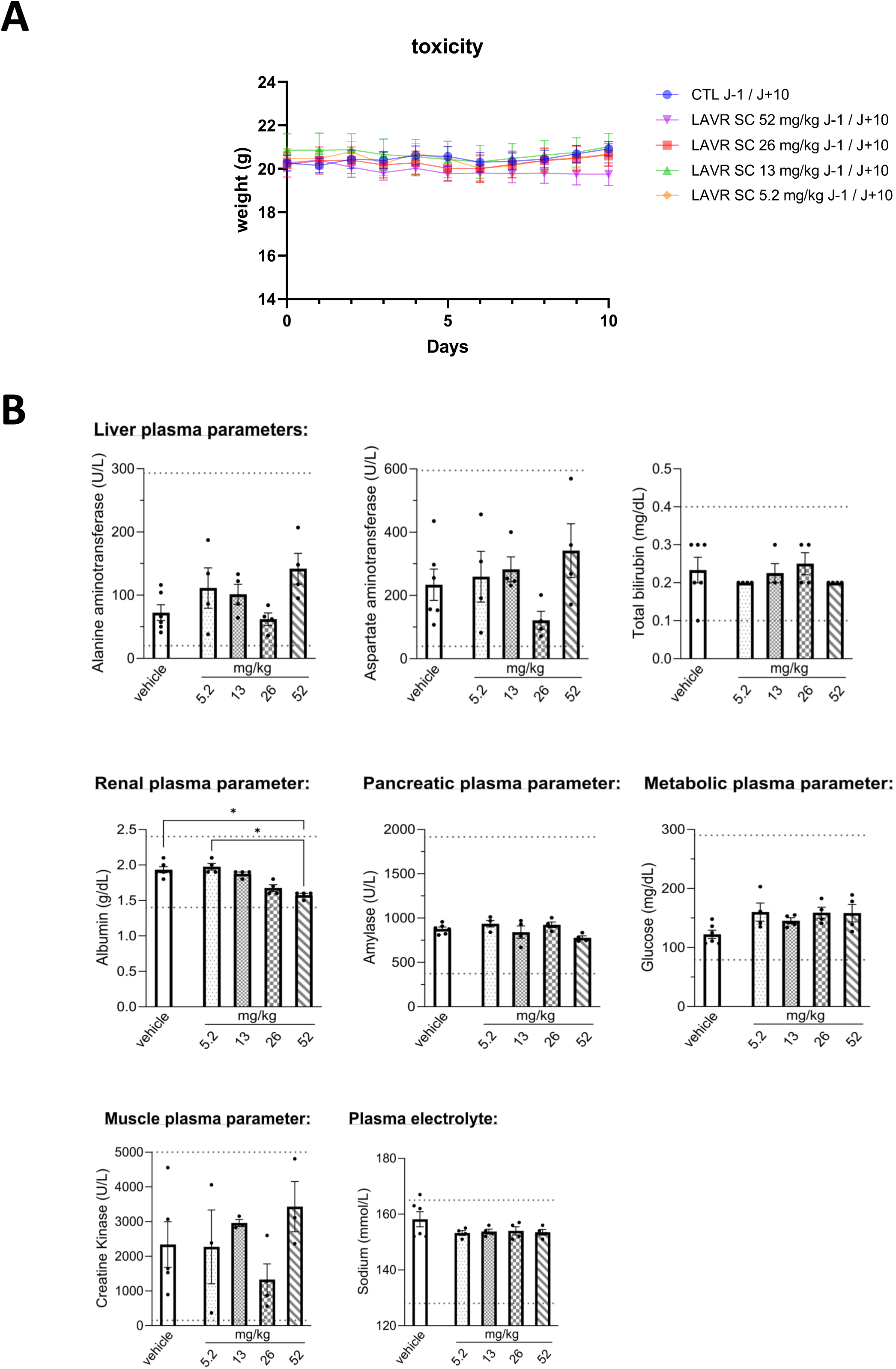
LAVR-289 is not toxic in mice. (A) Mice weight following daily SC injection of LAVR-289 for 10 days at the indicated concentrations. LAVR-289 is formulated in 10% DMSO / 10%Cremophor / 80% PBS. (B) Plasmatic parameters analyzed after LAVR-289 (5.2, 13, 26, or 52 mg/kg) or vehicle administration in distinct group of mice at day 10. Thresholds (min and max) for mice are indicated in grey dotted line. Kruskal-Wallis test followed by Dunn’s multiple comparisons test; * p < 0.05; n = 6 mice/group.

**Figure 5:**
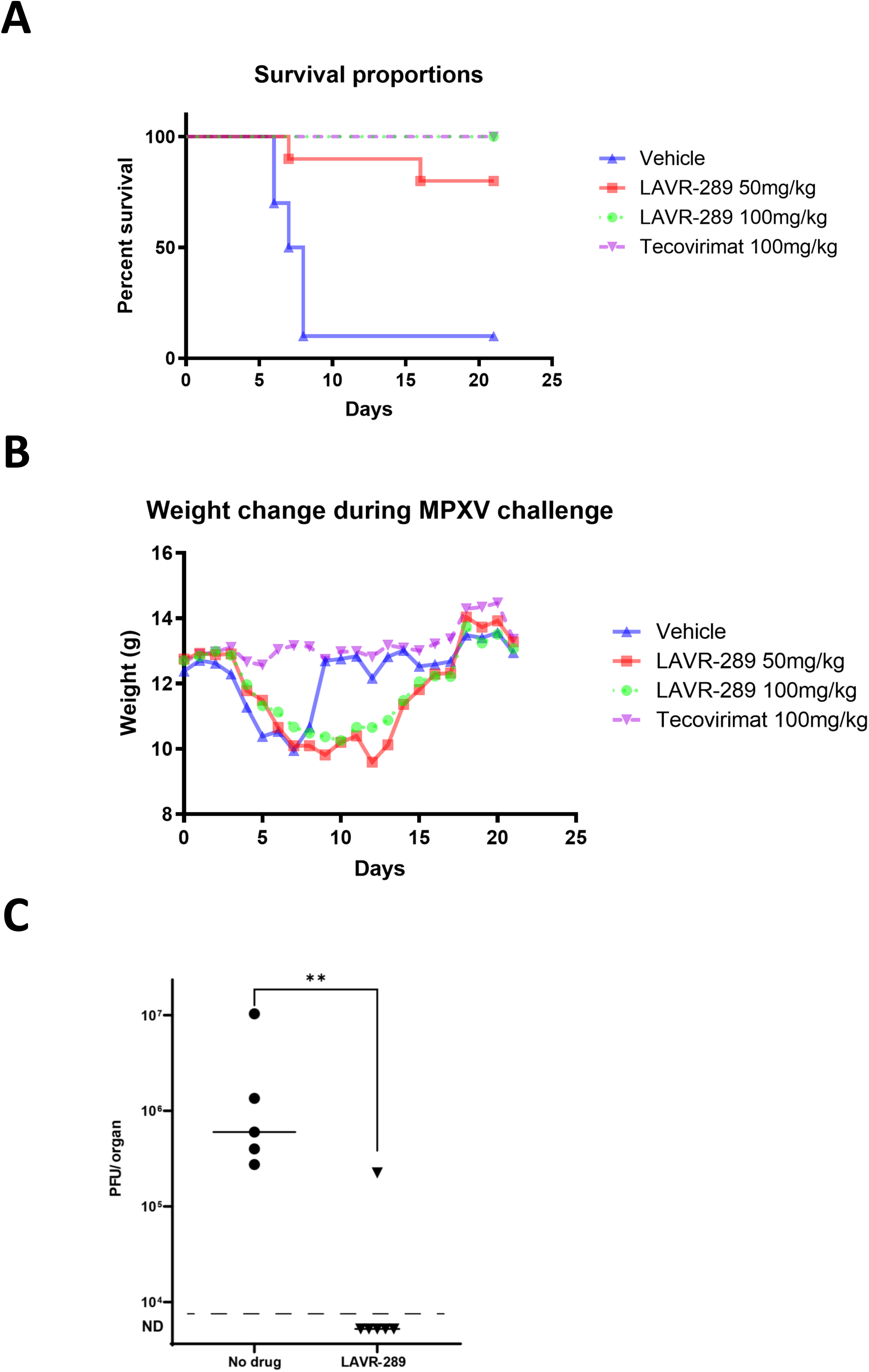
LAVR-289 reduces symptom severity and protects up to 100% of animals during a lethal MPXV challenge. (A) Probability of survival of the untreated and LAVR-289 groups during intranasal clade Ia MPXV challenge. Tecovirimat is used as a positive control. (B) Group weight curves during the LAVR-289 challenge. Tecovirimat treated mice at 100mg/kg do not lose weight. Infected control and LAVR-289 groups lose weight due to MPXV infection. (C) clade IIb MPXV viral burden in the lung of untreated groups or per os administration of 50mg/kg LAVR-289 starting 3 days post infection. Mice were sacrificed at day 7 post challenge. **: p<0.01 (Kolmogorov-Smirnov test)

### 3.6. LAVR-289 completely protects against lethality in a clade Ia MPXV challenge model and reduces viral lung burden in a clade IIb MPXV murine model

To investigate if LAVR-289 is able to inhibit MPXV induced mortality, we used CAST/EiJ mice (6-8 weeks old, 10 mice/group of LAVR-289) infected with clade Ia MPXV intranasally with 10^5^ PFU^27^. Tecovirimat was used as a positive control in this study (7 mice per group). As shown in Fig 6A, in untreated conditions, 9/10 animals succumbed to infection at Day 8. In stark contrast, 8/10 and 10/10 in the LAVR-289 treated animals survived the infection, as did the 7/7 tecovirimat treated animals, indicating that LAVR-289 treatment is able to prevent clade Ia MPXV-induced mortality when initiated one day post-infection. It is to be noted that a single animal in the 50mg/kg group died unexpectedly on Day 16 post exposure without any clinical signs of disease. Weights (Fig 6B) trends show that regardless of infection, LAVR-289 treated groups lose weight due to infection. Interestingly, the 100mg/kg seemed to mitigate weight loss. During the critical period of the disease (Days 7-9), when mortality in the control group occurs (clinical score of >8), LAVR-289 groups displayed lower clinical score, ranging between 4 and 5 (*data no shown*). Following this critical period, surviving mice tended to gain weight with decreasing clinical score reaching 0 by the end of the study. To expand upon these results, and to test whether LAVR-289 can impact the viral lung burden in infected animals, we used CAST/EiJ mice infected with a clade IIb MPXV intranasally with 10^4^ PFU either in untreated condition or using daily oral administration of LAVR-289 at 50mg/kg starting three days after infection. As shown in Fig. 6C, LAVR-289 was able to almost completely abolish viral replication in the lung of infected animals. Together, these results provide evidence that LAVR-289 can prevent MPXV induced mortality; possibly by a mechanism reducing virus replication in animals similar to evidence in the clade IIb MPXV murine model.

## 4. Discussion and summary

Nucleoside inhibitors are among the most represented class of antivirals approved on the market. More than 37.5% of all antivirals target the viral polymerase^28^. Currently approved antiviral that target poxviruses infection are limited, and only Brincidofovir and Tecovirimat are licensed by the FDA. These two treatments, one targeting virus replication (BCDV) and the other targeting viral egress (tecovirimat), quickly showed their limitations, in terms of efficacy, toxicity and rapid emergence of resistance^11,12,15,29^. There is therefore a constant need to develop new molecules against poxviruses having increased efficiency or a different part of the viral polymerase. Here, we have shown that LAVR-289 is a new acyclonucleoside phosphonate that displays nanomolar antiviral activities on a broad collection of poxviruses. Its antiviral activity is not restricted to poxvirus, as we detected broad spectrum antiviral activities on other dsDNA virus families^18^ [see Kappler-Gratias et al. and Quentin-Froignant et al., same issue]. LAVR-289 is able to completely prevent poxvirus replication *in vitro*. Treatment with LAVR-289 impacts formation of replication centers in infected cells, meaning that the compound acts early in the viral replication cycle and targets directly the viral DNA polymerase. Pulse chase experiment has shown that antiviral activity remains stable over time, indicating that the compound is stable in cell culture or that genome modification is a terminal phenotype and viral genome having incorporated the LAVR-289 cannot be repaired. LAVR-289 is active on all poxviruses tested so far and also active on patient samples (MPXV/France/IRBA2211/2022, CPXV CNR Orthopoxvirus, France) with maximum selectivity indexes reaching around 600. LAVR-289 also inhibits viruses of veterinary interest, such as MYXV, LSDV and AFSV where no antivirals or vaccines are available at large scale. Therefore, during an outbreak, LAVR-289 or similar compounds could be used to reduce the viral burden in an infected farm and in an effort to reduce spreading of the disease to surrounding farms. This may be helpful to reduce culling of herds.

One advantage of LAVR-289 is that it is an acyclic nucleoside phosphonate and serves as an analogue of a 5’-monophosphate nucleoside form; meaning that its initial (and rate-limiting) phosphorylation step by viral kinases is not required for its activity. This leads to a product being active on all TK- resistant strains [see Kappler gratias et al., same issue]. Mapping LAVR-289 resistance mutations highlighted the hydrophobic pocket of the insert 2 of the VACV viral DNA polymerase as the main target albeit we cannot exclude another domain of the polymerase to be involved. The C356Y single mutation is a known mutation conferring resistance to phosphonoacetic acid, a pyrophosphate analog, such as Foscarnet^26,30,31^. Therefore, it seems that phosphate binding pocket in the VACV polymerase contributes to LAVR-289 function. This is confirmed by the fact that HSV-1 Foscarnet resistant strain displays a 2.5-fold increase in LAVR-289 EC_50_ [Kappler gratias et al., same issue]. Remarkably, LAVR-289 is not toxic either when administered SC, IV or *per os* [see Quentin-Froignant, same issue], no modification of the weight of the animal or in the behavior could be observed. In MPXV challenge, we have shown that LAVR reduces clinical signs and increases survival of up to 100% of infected animals. LAVR-289 is also able to protect 100% hamsters infected with intravenous HAdvC6 when given *per os* (see Quentin-Froignant, same issue).

According to its innovative synthesis, broad spectrum antiviral activity, high stability in plasma, long half-life, absence of toxicity and its efficacy *in vivo*, this compound is a promising alternative to brincidofovir. In addition, the compound is extremely stable when in solution or in powder at -20 °C (24 months without loss of activity, *data not shown*), this is of special interest in case of stockpiling as a medical counter measure in case of poxvirus outbreak. As we saw the successive outbreaks of several viruses over the last years (i.e., SARS-CoV-2 and MPXV) development of new generation broad spectrum antivirals should be of high concern for the sake of general population.

## Acknowledgments

Authors thank French Defense Innovation Agency (RAPID program “Denalpovir” grant # 192906106), and Region Centre Val de Loire (APR-IR FINALS) for financial support, which made this study possible. ICOA UMR CNRS 7311 receives grants from the University of Orléans and from the CNRS as well as from FEDER (EX003677, EX011313, 2021-2027- 00022860), ANR Precyvir, Labex SYNORG (ANR-11-LABX-0029) and IRON (ANR-11-LABX-0018-01), and the SALSA platform for spectroscopic measurements. F.I. is supported by research grants from the *Service de Santé des Armées* and the *Direction Générale de l’Armement*. The clade IB MPXV animal studies was funded by the National Institute of Allergy and Infectious Diseases (NIAID), the National Institutes of Health (NIH), through an Interagency Agreement (# AAI24020-001-00000) with the U.S. Army Medical Research Institute (USAMRIID) to identify small-molecule orthopoxvirus inhibitors. The opinions, interpretations, conclusions, and recommendations presented are those of the author and are not necessarily endorsed by the U.S. Army or Department of Defense. Animal studies with clade Ia MPXV was conducted at USAMRIID under an Institutional Animal Care and Use Committee (IACUC) approved protocol in compliance with the Animal Welfare Act, PHS Policy, and other Federal statutes and regulations relating to animals and experiments involving animals. The facility where this research was conducted is accredited by the Association for Assessment and Accreditation of Laboratory Animal Care International and adheres to principles stated in the Guide for the Care and Use of Laboratory Animals, National Research Council, 2011. Authors thank Jean-Nicolas Tournier (IRBA and École du Val-de-Grâce, Paris, France) and François Caire-Maurisier (Pharmacie Centrale des Armées, Chanteau, France) for assistance. Authors thank Philippe Erbs (Transgene, Illkirch-Graffenstaden, France) for the use of TG6002 product and CREFRE unit for animal facility use.

## Author Contributions

Conceptualization: R.M, L.A.A., F.G. and F.I.; Experiments: E.M., E.Mo., G.F.V., S.K.G., L.B., S.T., T.M., C.C., P.F, T.G, P.B, M.K, D.G., A.M, A.A, Y.Y, K.T, G.A, V.R, J.S, K.S.W, S.D.J, G.W.J, R.G.P, E.M.M, R.M, F.I, FG ; Supervision: F.G. and F.I.; Writing – review & editing: F.G - R.M, F.G., M.K, L.A.A, E.M.M, K.T, G.A and F.I. Funding acquisition: L.A.A., F.G and F.I. All authors have read and agreed to the published version of the manuscript.

---

**Figure S1:**
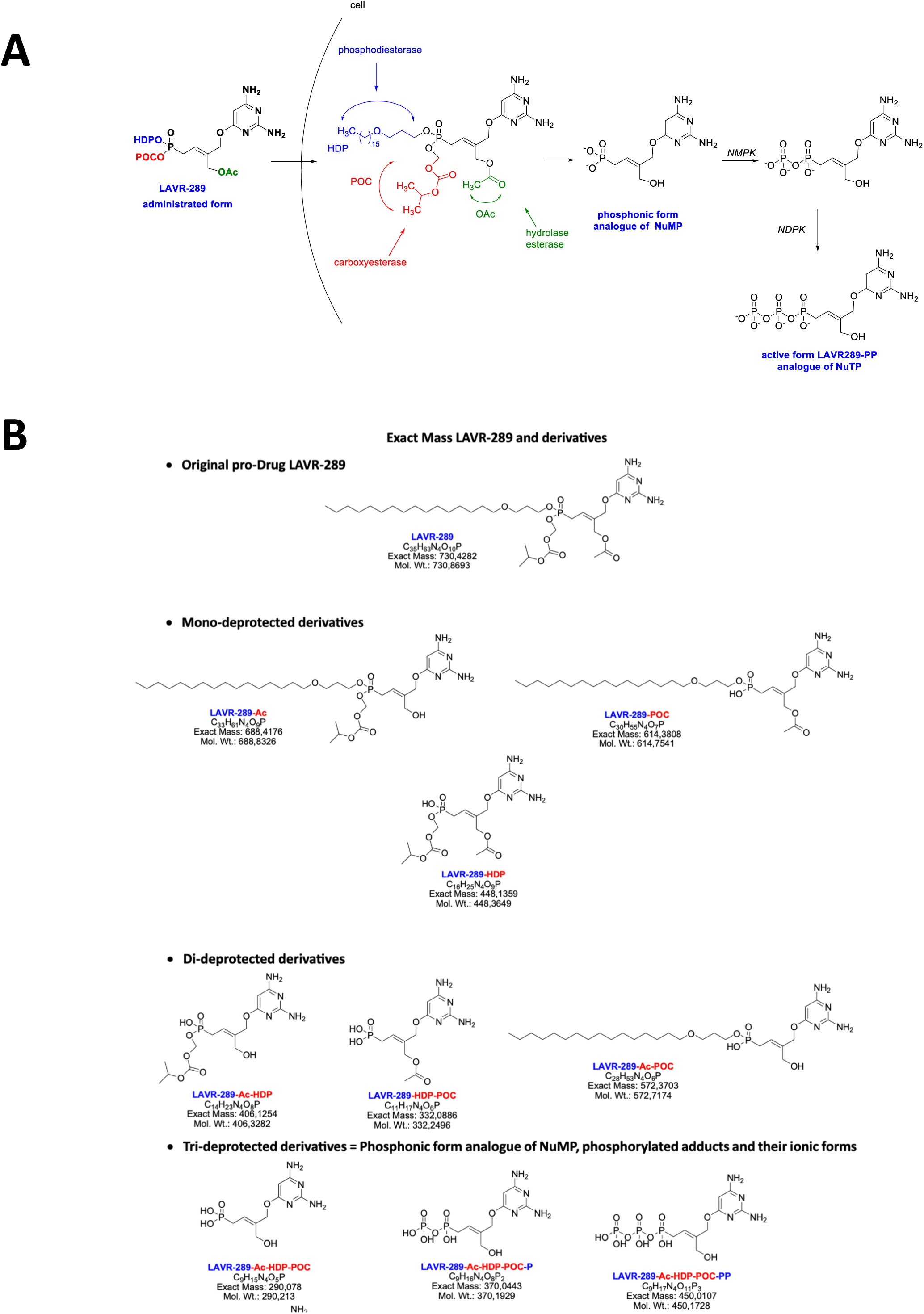
LAVR-289 proposed mechanism of action. (A) Putative mechanism of action of LAVR-289. Due to its prodrug moiety, LAVR-289 rapidly cross the cell envelope and is metabolized by enzymatic activities (phosphodiesterase, carboxylesterase and hydrolase esterase) to liberate the phosphonic acid which is successfully phosphorylated by nucleoside monophosphate kinase (NMPK) and nucleoside diphosphate kinase (NDPK) to give tri-P LAVR-289. Tri-P is then incorporated in replicating viral DNA and terminates replication. (B) All LAVR-289 putative mono, di and tri deprotected derivatives are shown. Removed biolabile group(s) on each metabolite are shown in red.

**Figure S2:**
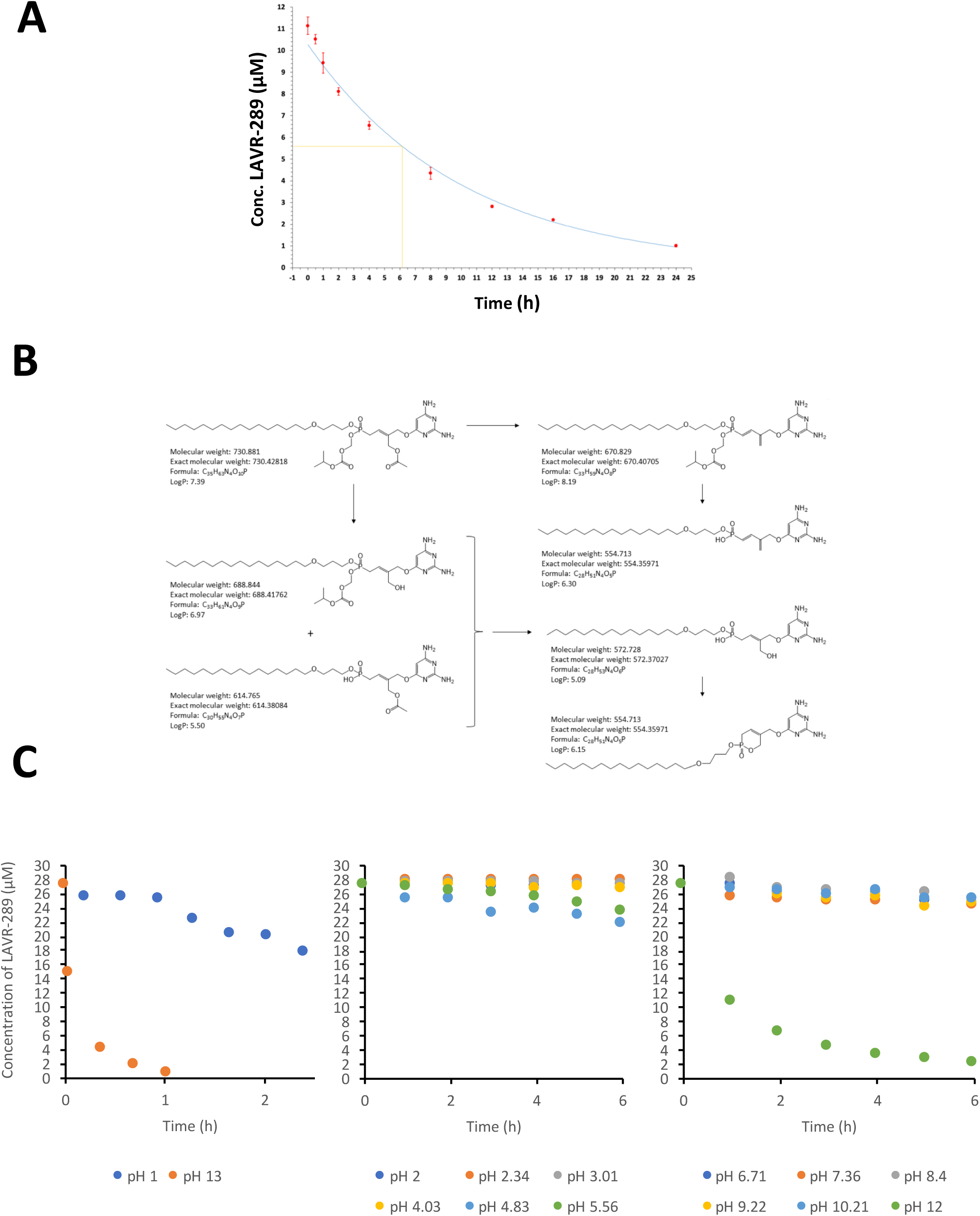
LAVR-289 stability in human plasma, detected metabolites and pH stability. (A) Profile of the stability kinetics of LAVR-289 in human plasma at 37°C (red points). Error bars are equal to twice of the residual standard deviation values calculated for each time (n = 3). Blue line represents the plot of the fitted 1^st^ order kinetics model found by nonlinear regression (r² = 0.9932). The half-life of the prodrug in the human plasma is estimated to be 6.16 h. (B) LAVR-289 degradation products detected in the human plasma by UPLC-HR-MS. LogP are calculated for neutral forms. (C) LAVR-289 was incubated in extreme pH (NaOH or HCl 0,1N left) or in universal buffer Britton-Robinson at intermediate pH. Stability of the LAVR-289 is assessed by HPLC at the indicated time.

**Figure S3:**
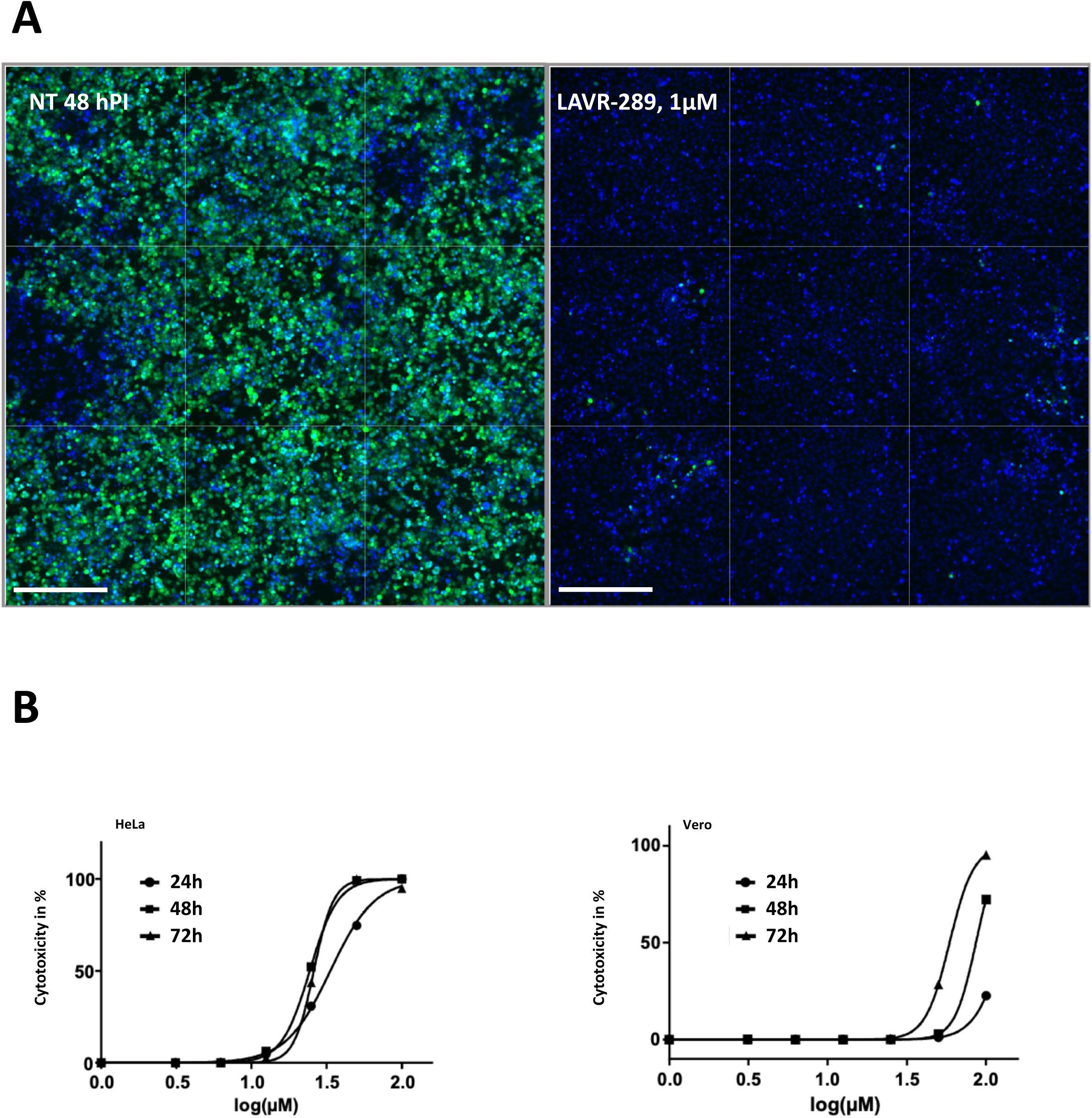
LAVR-289 impacts poxvirus replication without acute toxicity. (A)LAVR-289 impedes infection progression at high concentration. HeLa cells were infected with VACV ANCHOR at MOI 0.05 and incubated for 48 h. Cells were fixed and processed for HCS quantification. 9 fields reconstruction taken at 10X of untreated (left) and LAVR-289 1 µM treated cells (right) is shown (bar=400µm). (B) LAVR-289 toxicity assessed using the LDH cytotoxicity assay on HeLa cells (left) and Vero cells (right) following 1-, 2- or 3-days incubation. Calculated CC_50_ for Hela cells at Day 1, 2 and 3 are 33.46, 24.41 and 25.98 µM, respectively. For Vero cells, calculated CC_50_ at Day 1, 2 and 3 are 130.3, 86.22 and 58.79 µM, respectively.

**Figure S4:**
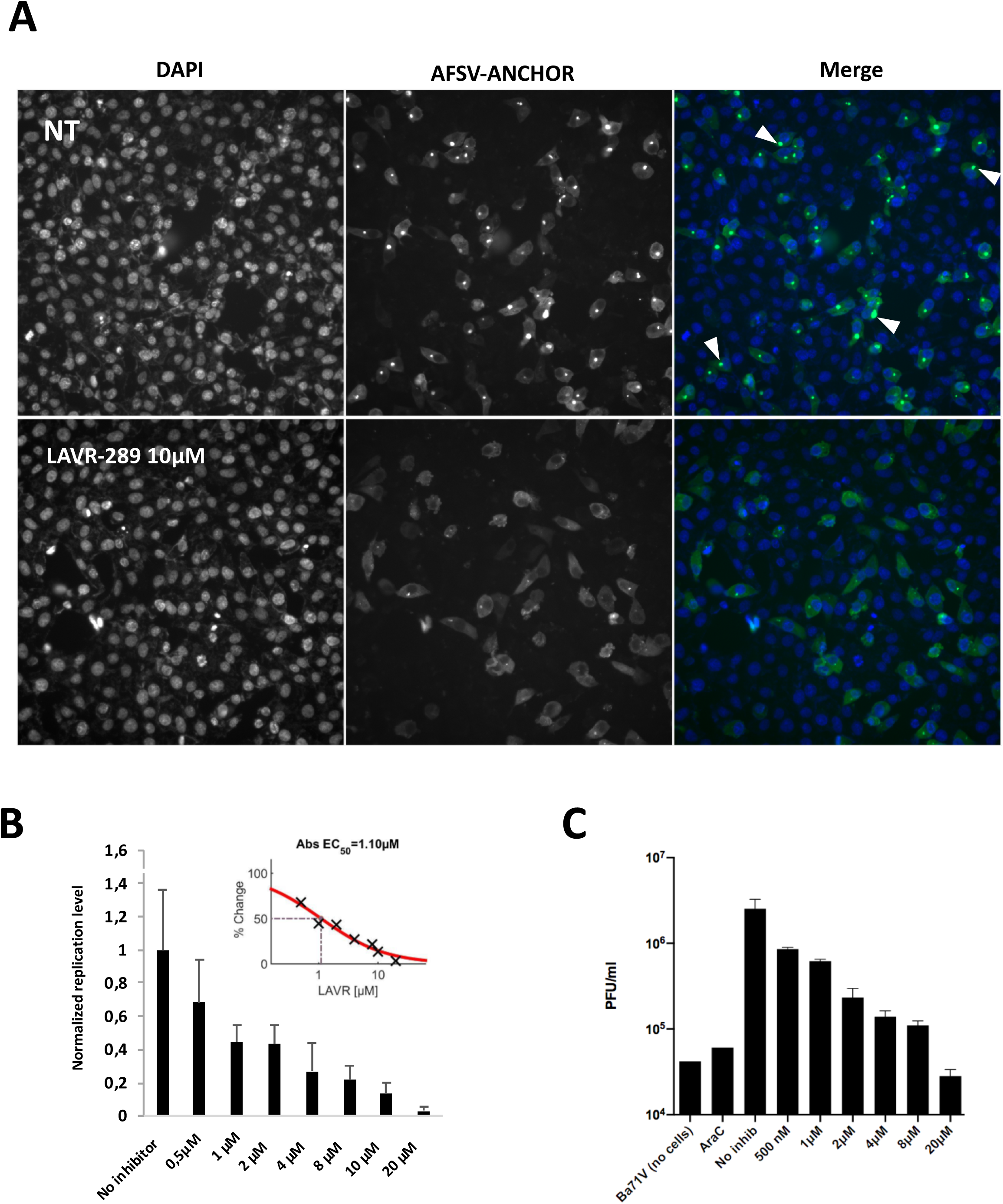
LAVR-289 inhibits AFSV replication measured by HCS or PFU quantification. (A) Single field representation of AFSV ANCHOR infection in the absence or presence of 10µM LAVR-289. Arrowheads show AFSV replication center. Note the strong reduction of replication centers surface and intensity in LAVR-289 treated cells (bar=400µm) (C) Measurement of replication rate using compartmental analysis coupled to spot detector and associated calculation of EC_50_ (1.1µM). (D) Quantification (in PFU/mL) of AFSV BA71V ANCHOR produced in Vero cells treated with increasing concentration of LAVR-289. Commercially available AraC (1-β-D-arabinosyl-cytosine) was used as positive control.

## References

1. Moss, B.; Smith, G., L. Poxviridae: The Viruses and Their Replication. In Fields Virology; Wolters Klumer, 2021; Vol. 2.

2. Smallpox Eradication: Destruction of Variola Virus Stocks (Https://Iris.Who.Int/Bitstream/Handle/10665/79354/E10.Pdf?Sequence=1), 1999. https://iris.who.int/bitstream/handle/10665/79354/e10.pdf?sequence=1.

(3) Berche, P. Life and Death of Smallpox. Presse Medicale Paris Fr. 1983 2022, 51 (3), 104117. 10.1016/j.lpm.2022.104117.

(4) Bunge, E. M.; Hoet, B.; Chen, L.; Lienert, F.; Weidenthaler, H.; Baer, L. R.; Steffen, R. The Changing Epidemiology of Human Monkeypox-A Potential Threat? A Systematic Review. PLoS Negl. Trop. Dis. 2022, 16 (2), e0010141. 10.1371/journal.pntd.0010141.

5. WHO (World Health Organization). MpoxC: Global Strategic Preparedness and Response Plan, 2024. https://cdn.who.int/media/docs/default-source/documents/health-topics/monkeypox/jmo_who_sprp-mpox_2024_final_digital.pdf?sfvrsn=3a670f76_1&download=true.

(6) Wolff Sagy, Y.; Zucker, R.; Hammerman, A.; Markovits, H.; Arieh, N. G.; Abu Ahmad, W.; Battat, E.; Ramot, N.; Carmeli, G.; Mark-Amir, A.; Wagner-Kolasko, G.; Duskin-Bitan, H.; Yaron, S.; Peretz, A.; Arbel, R.; Lavie, G.; Netzer, D. Real-World Effectiveness of a Single Dose of Mpox Vaccine in Males. Nat. Med. 2023, 29 (3), 748–752. 10.1038/s41591-023-02229-3.

(7) Delaune, D.; Iseni, F. Drug Development against Smallpox: Present and Future. Antimicrob. Agents Chemother. 2020, 64 (4), e01683–19. 10.1128/AAC.01683-19.

(8) Jordan, R.; Leeds, J. M.; Tyavanagimatt, S.; Hruby, D. E. Development of ST-246® for Treatment of Poxvirus Infections. Viruses 2010, 2 (11), 2409–2435. 10.3390/v2112409.

9. Frenois-Veyrat G; Gallardo F; Gorgé O; Marcheteau E; Ferraris O; Baidaliuk A; Favier Al; Enfroy C; Holy X; Lourenco J; Khoury R; Nolent F; Grosenbach Dw; Hruby De; Ferrier A; Iseni F; Simon-Loriere E; Tournier Jn. Tecovirimat Is Effective against Human Monkeypox Virus in Vitro at Nanomolar Concentrations. Nat. Microbiol. 2022, 7 (12). 10.1038/s41564-022-01269-8.

10. The antiviral tecovirimat is safe but did not improve clade I mpox resolution in Democratic Republic of the Congo. National Institutes of Health (NIH). https://www.nih.gov/news-events/news-releases/antiviral-tecovirimat-safe-did-not-improve-clade-i-mpox-resolution-democratic-republic-congo (accessed 2025-01-20).

11. NIH Study Finds Tecovirimat Was Safe but Did Not Improve Mpox Resolution or Pain. National Institutes of Health (NIH). https://www.nih.gov/news-events/news-releases/nih-study-finds-tecovirimat-was-safe-did-not-improve-mpox-resolution-or-pain (accessed 2025-01-20).

(12) Garrigues, J. M.; Hemarajata, P.; Karan, A.; Shah, N. K.; Alarcón, J.; Marutani, A. N.; Finn, L.; Smith, T. G.; Gigante, C. M.; Davidson, W.; Wynn, N. T.; Hutson, C. L.; Kim, M.; Terashita, D.; Balter, S. E.; Green, N. M. Identification of Tecovirimat Resistance-Associated Mutations in Human Monkeypox Virus - Los Angeles County. Antimicrob. Agents Chemother. 2023, e0056823. 10.1128/aac.00568-23.

(13) Alarcón, J.; Kim, M.; Terashita, D.; Davar, K.; Garrigues, J. M.; Guccione, J. P.; Evans, M. G.; Hemarajata, P.; Wald-Dickler, N.; Holtom, P.; Garcia Tome, R.; Anyanwu, L.; Shah, N. K.; Miller, M.; Smith, T.; Matheny, A.; Davidson, W.; Hutson, C. L.; Lucas, J.; Ukpo, O. C.; Green, N. M.; Balter, S. E. An Mpox-Related Death in the United States. N. Engl. J. Med. 2023, 388 (13), 1246– 1247. 10.1056/NEJMc2214921.

(14) De Clercq, E.; Holý, A.; Rosenberg, I.; Sakuma, T.; Balzarini, J.; Maudgal, P. C. A Novel Selective Broad-Spectrum Anti-DNA Virus Agent. Nature 1986, 323 (6087), 464–467. 10.1038/323464a0.

(15) Adler, H.; Gould, S.; Hine, P.; Snell, L. B.; Wong, W.; Houlihan, C. F.; Osborne, J. C.; Rampling, T.; Beadsworth, M. B.; Duncan, C. J.; Dunning, J.; Fletcher, T. E.; Hunter, E. R.; Jacobs, M.; Khoo, S. H.; Newsholme, W.; Porter, D.; Porter, R. J.; Ratcliffe, L.; Schmid, M. L.; Semple, M. G.; Tunbridge, A. J.; Wingfield, T.; Price, N. M.; NHS England High Consequence Infectious Diseases (Airborne) Network. Clinical Features and Management of Human Monkeypox: A Retrospective Observational Study in the UK. Lancet Infect. Dis. 2022, 22 (8), 1153–1162. 10.1016/S1473-3099(22)00228-6.

(16) Ruedas-Torres, I.; Thi to Nga, B.; Salguero, F. J. Pathogenicity and Virulence of African Swine Fever Virus. Virulence 15 (1), 2375550. 10.1080/21505594.2024.2375550.

(17) Guo, S.; Zhang, Y.; Liu, Z.; Wang, D.; Liu, H.; Li, L.; Chen, Q.; Yang, D.; Liu, Q.; Guo, H.; Mou, S.; Chen, H.; Wang, X. Brincidofovir Is a Robust Replication Inhibitor against African Swine Fever Virus in Vivo and in Vitro. Emerg. Microbes Infect. 2023, 12 (2), 2220572. 10.1080/22221751.2023.2220572.

(18) Bessières, M.; Roy, V.; Abuduani, T.; Favetta, P.; Snoeck, R.; Andrei, G.; Moffat, J.; Gallardo, F.; Agrofoglio, L. A. Synthesis of LAVR-289, a New [(Z)-3-(Acetoxymethyl)-4-(2,4-Diaminopyrimidin- 6-Yl)Oxy-but-2-Enyl]Phosphonic Acid Prodrug with Pronounced Antiviral Activity against DNA Viruses. Eur. J. Med. Chem. 2024, 271, 116412. 10.1016/j.ejmech.2024.116412.

(19) Gallardo, F.; Schmitt, D.; Brandely, R.; Brua, C.; Silvestre, N.; Findeli, A.; Foloppe, J.; Top, S.; Kappler-Gratias, S.; Quentin-Froignant, C.; Morin, R.; Lagarde, J.-M.; Bystricky, K.; Bertagnoli, S.; Erbs, P. Fluorescent Tagged Vaccinia Virus Genome Allows Rapid and Efficient Measurement of Oncolytic Potential and Discovery of Oncolytic Modulators. Biomedicines 2020, 8 (12), E543. 10.3390/biomedicines8120543.

(20) Mariamé, B.; Kappler-Gratias, S.; Kappler, M.; Balor, S.; Gallardo, F.; Bystricky, K. Real-Time Visualization and Quantification of Human Cytomegalovirus Replication in Living Cells Using the ANCHOR DNA Labeling Technology. J. Virol. 2018, 92 (18), e00571–18. 10.1128/JVI.00571-18.

(21) Kappler-Gratias, S.; Bucher, L.; Desbois, N.; Rousselin, Y.; Bystricky, K.; Gros, C. P.; Gallardo, F. A3- and A2B-Fluorocorroles: Synthesis, X-Ray Characterization and Antiviral Activity Evaluation against Human Cytomegalovirus Infection. RSC Med. Chem. 2020, 11 (7), 783–801. 10.1039/d0md00127a.

(22) Boutin, L.; Mosca, E.; Iseni, F. Efficient Method for Generating Point Mutations in the Vaccinia Virus Genome Using CRISPR/Cas9. Viruses 2022, 14 (7), 1559. 10.3390/v14071559.

(23) Martin, V.; Guerra, B.; Hernaez, B.; Kappler-Gratias, S.; Gallardo, F.; Guerra, M.; Andres, G.; Alejo, A. A Novel Live DNA Tagging System for African Swine Fever Virus Shows That Bisbenzimide Hoechst 33342 Can Effectively Block Its Replication. Antiviral Res. 2024, 230, 105973. 10.1016/j.antiviral.2024.105973.

(24) Kuai, L.; Sun, J.; Peng, Q.; Zhao, X.; Yuan, B.; Liu, S.; Bi, Y.; Shi, Y. Cryo-EM Structure of DNA Polymerase of African Swine Fever Virus. Nucleic Acids Res. 2024, 52 (17), 10717–10729. 10.1093/nar/gkae739.

(25) Sèle, C.; Gabel, F.; Gutsche, I.; Ivanov, I.; Burmeister, W. P.; Iseni, F.; Tarbouriech, N. Low-Resolution Structure of Vaccinia Virus DNA Replication Machinery. J. Virol. 2013, 87 (3), 1679– 1689. 10.1128/JVI.01533-12.

(26) Tarbouriech, N.; Ducournau, C.; Hutin, S.; Mas, P. J.; Man, P.; Forest, E.; Hart, D. J.; Peyrefitte, C. N.; Burmeister, W. P.; Iseni, F. The Vaccinia Virus DNA Polymerase Structure Provides Insights into the Mode of Processivity Factor Binding. Nat. Commun. 2017, 8 (1), 1455. 10.1038/s41467-017-01542-z.

(27) Americo, J. L.; Moss, B.; Earl, P. L. Identification of Wild-Derived Inbred Mouse Strains Highly Susceptible to Monkeypox Virus Infection for Use as Small Animal Models. J. Virol. 2010, 84 (16), 8172–8180. 10.1128/JVI.00621-10.

(28) Chaudhuri, S.; Symons, J. A.; Deval, J. Innovation and Trends in the Development and Approval of Antiviral Medicines: 1987–2017 and Beyond. Antiviral Res. 2018, 155, 76–88. 10.1016/j.antiviral.2018.05.005.

(29) The antiviral tecovirimat is safe but did not improve clade I mpox resolution in Democratic Republic of the Congo. National Institutes of Health (NIH). https://www.nih.gov/news-events/news-releases/antiviral-tecovirimat-safe-did-not-improve-clade-i-mpox-resolution-democratic-republic-congo (accessed 2024-12-09).

(30) Andrei, G.; Gammon, D. B.; Fiten, P.; De Clercq, E.; Opdenakker, G.; Snoeck, R.; Evans, D. H. Cidofovir Resistance in Vaccinia Virus Is Linked to Diminished Virulence in Mice. J. Virol. 2006, 80 (19), 9391–9401. 10.1128/JVI.00605-06.

(31) Taddie, J. A.; Traktman, P. Genetic Characterization of the Vaccinia Virus DNA Polymerase: Cytosine Arabinoside Resistance Requires a Variable Lesion Conferring Phosphonoacetate Resistance in Conjunction with an Invariant Mutation Localized to the 3’-5’ Exonuclease Domain. J. Virol. 1993, 67 (7), 4323–4336.

